# Autism-associated PTCHD1 missense variants bind to the SNARE-associated protein, SNAPIN, but exhibit impaired subcellular trafficking

**DOI:** 10.1101/2024.02.29.582618

**Authors:** Stephen F. Pastore, Connie T.Y. Xie, RoyaDerwish, Tahir Muhammad, Tereza Blahova, Sierra C. El-masri, Paul W. Frankland, Paul A. Hamel, John B. Vincent

## Abstract

**Background:** Patched domain-containing 1 (*PTCHD1*) is a susceptibility gene for autism spectrum disorder and intellectual disability. Its function in brain development and neurotransmission remains elusive. Studies have sought to characterize PTCHD1 function by elucidating its neural network of interacting proteins. However, given the current paucity of functional information, many PTCHD1 missense variants in clinical databases are classified as variants of uncertain significance (VUSs), severely limiting the healthcare resources available to patients and families.

**Methods:** A yeast two-hybrid assay was used to identify synaptic PTCHD1-interacting proteins. Candidate binding partners were validated by cloning; transient over-expression in HEK293T cells, followed by co-immunoprecipitation and immunoblotting; and immunocytochemistry in differentiated P19 cells. To evaluate the pathogenicity of clinical missense variants, site-directed mutagenesis was employed, followed by transient over-expression and immunocytochemistry in non-neuronal (HEK293T) and neuronal (Neuro-2A cells) systems.

**Results:** A novel interaction was identified between the first lumenal loop of PTCHD1 and the SNARE-associated protein SNAPIN, which is implicated in synaptic vesicle exocytosis. Clinically associated missense variants within this region did not disrupt SNAPIN binding, indicating that the pathoetiology of these variants is unrelated to this interaction. However, six of the 12 missense variants tested exhibited pronounced retention within the endoplasmic reticulum, and impaired neuronal and non-neuronal trafficking to the plasma membrane.

**Conclusions:** These data yield insights regarding the role of PTCHD1 in neurodevelopment and neurotransmission, and suggest a neuropathological mechanism for missense variants. These findings provide a platform for diagnostic assay and VUS interpretation, allowing for clinical re-classification of these variants.

## Introduction

Autism spectrum disorder (ASD) is a heterogeneous neurodevelopmental disorder typified by social and communicative deficiencies, restricted interests, and repetitive sensory-motor behaviours (1). ASD exhibits 50-80% comorbidity with intellectual disability (ID) (2). Genetics appear to strongly influence the pathoetiology of ASD, with a recent meta-analysis concluding that 74-93% of ASD risk is heritable (3). Genome-wide studies of affected cohorts have identified numerous ASD and ID susceptibility genes. *PTCHD1*, on Xp22.11, was first implicated in the pathoetiology of ASD and ID in 2008 (4). Numerous highly-penetrant rare genomic deletion variants and loss-of-function coding variants have been subsequently detected in *PTCHD1* (5), and many rare missense variants have been identified through clinical diagnostics (for example, see ClinVar: www.ncbi.nlm.nih.gov/clinvar).

*In silico* analyses have predicted PTCHD1 to be a multi-pass transmembrane protein consisting of 12 transmembrane domains (TMDs). Exogenous PTCHD1 has been found to localize within cell membranes (6–8), and localizes to the post-synaptic density (PSD) dendritic spines of neurons (7). In addition, PTCHD1 is predicted to possess two distinct sterol sensing domain-like modules, two large lumenal loops, and a C-terminal PDZ-binding domain [reviewed in (5)]. Functionally, a definitive cellular role for PTCHD1 has yet to be elucidated. PTCHD1 exhibits 21.17% sequence homology with the cholesterol-trafficking transmembrane protein Niemann-Pick disease, type C1 (5), and has correspondingly been reported to directly bind cholesterol (9).. To delineate a putative function of PTCHD1, previous research has sought to characterize its network of interacting proteins in neurons. These studies have employed affinity purification to identify and validate several putative binding partners of PTCHD1 *in vitro*. In this regard, both PSD protein 95 (PSD95) and Synapse-Associated Protein 102 (SAP102) have been reported to interact with PTCHD1 via its C-terminal PDZ-binding domain (7,10).

Over 300 rare missense variants have been identified in *PTCHD1* through clinical diagnostics in autism and ID individuals (11,12). However, molecular mechanisms of pathology exist for a very small subset of these variants (8,13,14), and the vast majority are interpreted as variants of uncertain significance (VUSs) due to the lack of supporting segregation or functional data. This diagnostic uncertainty negatively impacts the integrative healthcare options available to many families. Functional evidence from molecular studies that would enable re-classification of PTCHD1 variants from VUS to ‘likely pathogenic’ would provide an explanation for individual challenges and enable thousands of families to access additional healthcare resources, including testing of carrier status in female relatives and appropriate genetic counselling.

The objective of the present study was to agnostically identify additional synaptic proteins that can interact with PTCHD1, beyond interactions through of its C-terminal PDZ-binding domain, in order to elaborate on its potential function(s) in neurons. Furthermore, we sought to evaluate PTCHD1 VUSs in both neuronal and non-neuronal systems, in order to provide evidence for their possible clinical re-classification.

## Methods and Materials

### Yeast Two-Hybrid Screen

The Matchmaker Gold Yeast Two-Hybrid System (Takara Bio; Kusatsu, Japan) was used. Briefly, lumenal loop 1 (amino acids p.Glu49-p.Arg270) or a fusion of lumenal loop 1 to lumenal loop 2 (amino acids p.Gln521-p.Ser695) of the human ortholog of *PTCHD1* was cloned into the pGBKT7 vector in-frame with the DNA-binding domain of the Gal4 transcription factor. Constructs were separately transformed into the *S. cerevisiae* strain Y2HGold and independently used as bait to probe two separate cDNA libraries fused to the activation domain of Gal4: 1) Mate & Plate Library Mouse Embryo Day 11, and 2) Normalized Mate & Plate Library Adult Human Brain (Takara Bio; Kusatsu, Japan). Three independent screens were performed for each library. Positive clones were selected on synthetic dropout medium in the absence of tryptophan, leucine, histidine, and adenine, and supplemented with X-gal. Positive clones were subjected to plasmid DNA extraction and transformation into *E. coli*, and plasmids from individual colonies were then amplified by PCR and Sanger sequencing was used to identify the insert.

### Cloning

The complete coding sequences of the mouse orthologs of *Ptchd1* and the SNARE-associated protein *Snapin*, which exhibit 98.1% and 97.8% protein sequence homology, respectively, with their human orthologs, were separately cloned into the expression vector pcDNA3.1 myc-His B (Thermo Fisher Scientific; Waltham, MA). *Snapin* was cloned in-frame with the downstream myc-His tags. For *Ptchd1*, the stop codon was included, and a 3xFlag epitope tag was subsequently inserted at the N-terminus. Finally, the coding sequences of 3xFlag-tagged *Ptchd1* lumenal loop 1 (amino acids p.Val48-p.Arg266), lumenal loop 2 (amino acid p.Tyr499-p.Ala698), and a lumenal loop 1-loop 2 chimeric protein were cloned into pcDNA3.1 myc-His B, succeeded by a stop codon. The human ortholog of *PTCHD1* was recombined into the vector pDEST53 (Thermo Fisher Scientific) in-frame with GFP via Gateway cloning, and human SNAPIN cloned into pcDNA3.1 myc-His B. Primer sequences used for expression cloning are provided in Table 1.

**Table 1.**
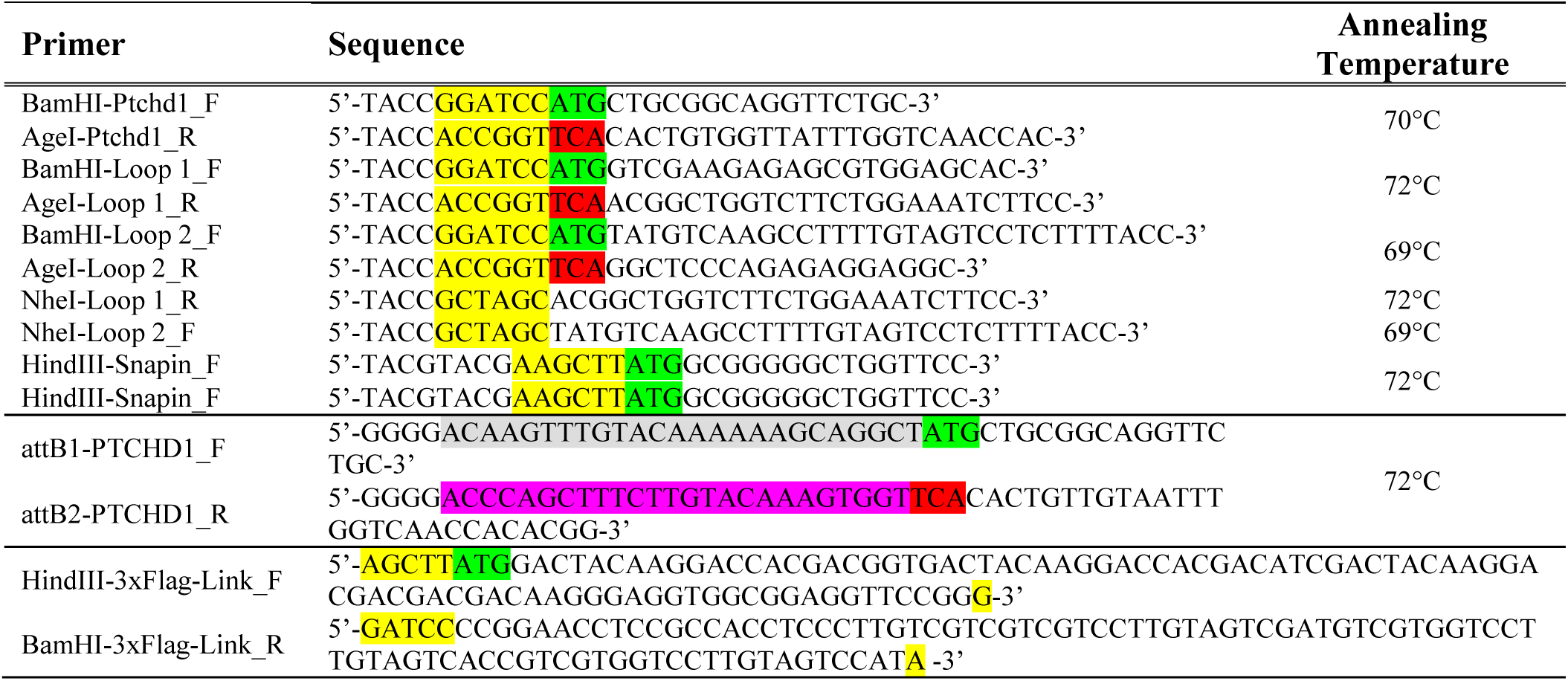
Oligonucleotides used to generate expression constructs. Full or partial restriction sites, AUG codons, and stop codons are highlighted in yellow, green, and red, respectively. For Gateway cloning, attB1 and attB2 sequences are denoted in grey and magenta, respectively. Annealing temperatures for primer pairs are also shown.

Site-directed mutagenesis was employed to generate the clinically-reported *Ptchd1* missense variants evaluated in this study (Table 2). The amino acid residues for all 14 missense variants evaluated are conserved between human and mouse (see Supplementary Figure S5). Briefly, the wildtype *3xFlag-Ptchd1-pcDNA3.1* vector was amplified using Q5 high-fidelity DNA polymerase (NEB; Ipswich, MA), with the forward primer containing the mutant codon of interest. Next, purified PCR products were phosphorylated and re-circularized with DNA ligase. All mutant constructs were confirmed by Sanger sequencing (Supplemental figures S1 and S2). Primer sequences used for site-directed mutagenesis are provided in Table 3.

**Table 2.**
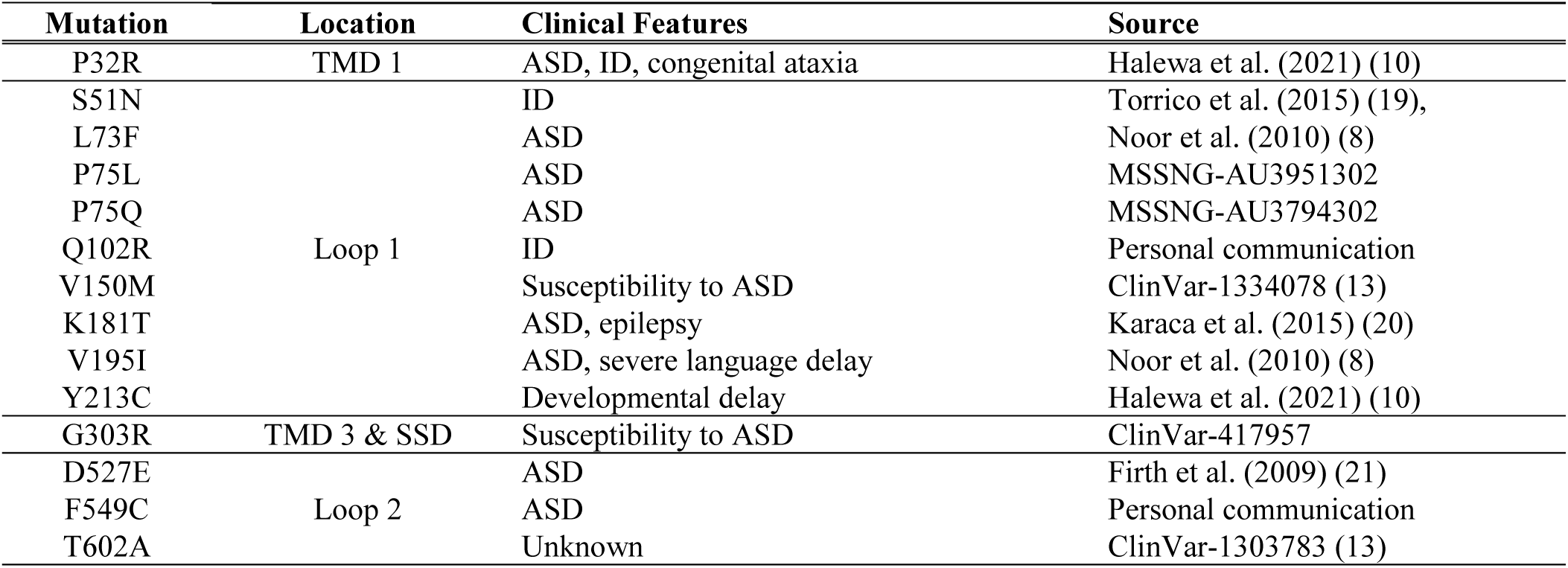
Location and clinical information for PTCHD1 missense variants. Five loop 1 variants were selected, three loop 2 variants, plus one variant in **t**ransmembrane domain (TMD) 1, and 1 in TMD3 that is also within the sterol sensing domain (SSD).

**Table 3.**
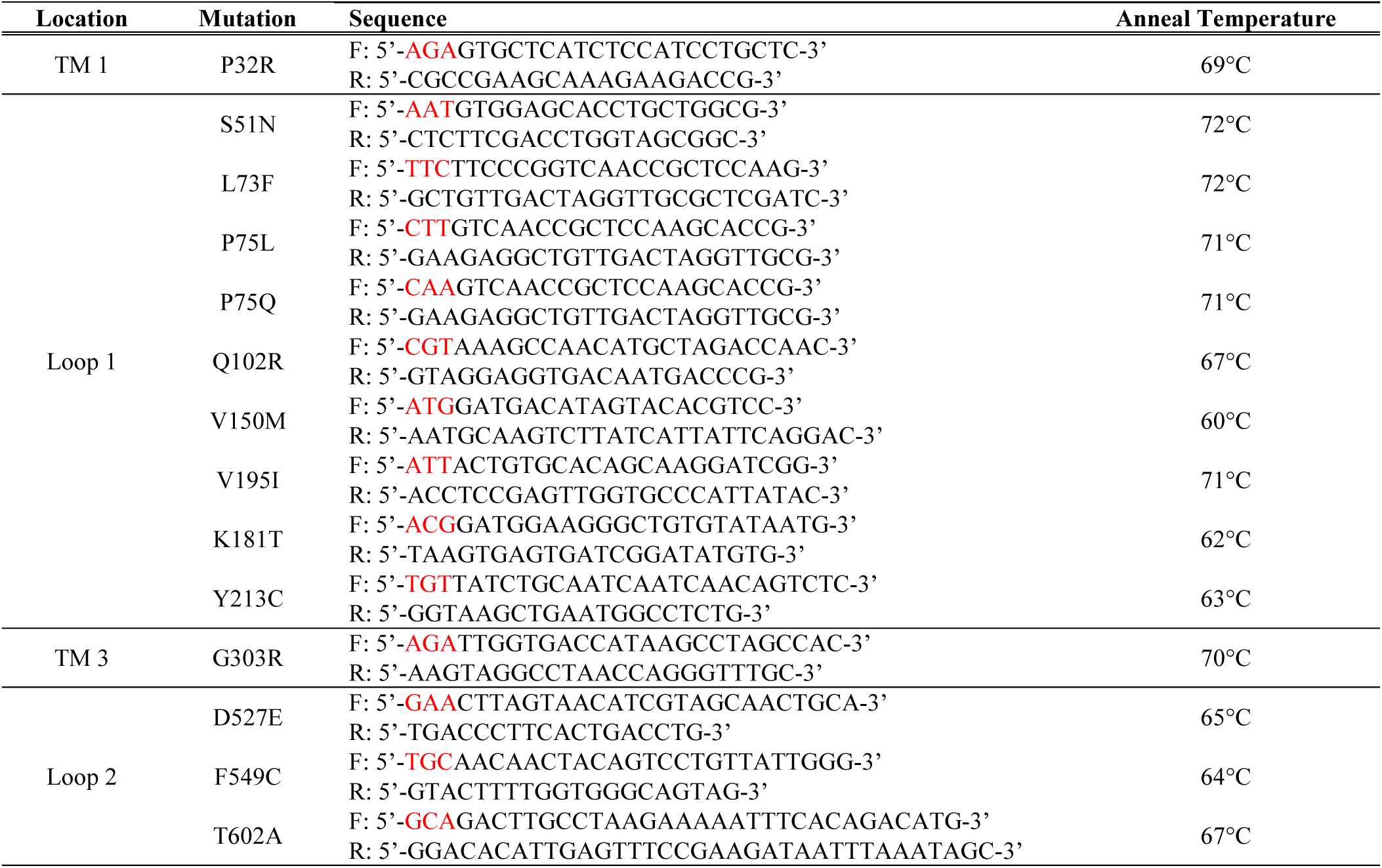
Oligonucleotides used for site-directed mutagenesis to generate missense variants in *Ptchd1*. Mutant codons are shown in red. Annealing temperatures for primer pairs are also shown.

### Cell Culture and Neuronal Differentiation

HEK293T cells (ATCC; CRL-3216) were maintained in DMEM supplemented with 10% FBS (Wisent; Saint-Jean-Baptiste, QC, Canada) and 1% PS (Wisent). Cells were passaged every 48-72 hours at semi-confluence by trypsinization (Wisent). P19 mouse embryonal carcinoma cells (ATCC; CRL-1825) were maintained in α-MEM supplemented with 7.5% newborn calf serum (Thermo Fisher Scientific), 2.5% FBS, and 1% PS. Undifferentiated cells were passaged every 48-72 hours at semi-confluence. For immunocytochemical staining, 24 hours prior to neuronal differentiation, Lipofectamine 3000 (Thermo Fisher Scientific) was utilized according to the manufacturer’s instructions to co-transfect *GFP-Ptchd1* and *Snapin-myc* into the cells. To induce neuronal differentiation, undifferentiated cells were grown on non-adherent bacterial-grade plates (Thermo Fisher Scientific) supplemented with 1 μM all-*trans* RA (Cell Signaling) for four days. On the fourth day, aggregates were plated onto sterile coverslips in 12-well plates, and allowed to differentiate for six days. Mouse Neuro-2a (N2a) neuroblastoma cells were seeded at a density of 70% confluency on coverslips in 6-well plates. They were cultured in Dulbecco’s Modified Eagle Medium (DMEM) with high glucose (4.5 g/L) supplemented with 10% (*v*/*v*) fetal bovine serum (FBS, Gibco) and containing 1% (*v*/*v*) antibiotics (100 U/mL penicillin, 100 mg/mL streptomycin; Sigma-Aldrich, St Louis, MO). Differentiation of N2A cells was induced by switching media to 4% FBS and adding retinoic acid to a final concentration of 2µM. For transfections, DNA constructs were mixed with 0.1% polyethylenimine (PEI; 1µgDNA:3µl PEI) (15) and added to N2A cells.

### Co-Immunoprecipitation

HEK293T cells were re-seeded in 6-well culture plates. 24 hours later, Lipofectamine 3000 (Thermo Fisher Scientific) was utilized according to the manufacturer’s instructions to co-transfect 2 μg of *3xFlag-Ptchd1* and 500 ng of *Snapin-myc* expression plasmids. 48 hours later, cells were washed with ice-cold PBS and lysed in immunoprecipitation (IP) buffer (50 mM Tris-HCl, pH 8.0, 150 mM NaCl, 1% NP-40) supplemented with cOmplete™ Mini EDTA-free protease inhibitors (MilliporeSigma, Burlington, MA). Lysates were incubated for 10 minutes on ice, followed by centrifugation at 10,000 x *g* for 10 minutes at 4°C. Supernatants were quantified using the Bradford total protein assay (Bio-Rad; Hercules, CA). For co-IP experiments, 500 μg of lysates were incubated with 1 μg of mouse α-Flag M2 antibody (Sigma-Aldrich; St. Louis, MO) in a final volume of 1 mL of IP buffer overnight at 4°C with rotation. The next day, lysates were incubated with 50 μl of Dynabeads Protein G (Thermo Fisher Scientific) for two hours at room temperature with rotation. Antibody-protein complexes were then separated magnetically, washed three times with PBS, and eluted from the Dynabeads using elution buffer (75 mM Glycine-HCl, pH 2.7). Co-transfected lysates were all immunoprecipitated using a mouse anti-Flag antibody, and then subsequent immunoblots were probed for Flag using a rabbit anti-Flag antibody, as indicated in Table 4. Two biological replicates were performed for co-IP experiments.

**Table 4.** Antibodies and dilutions used for IP, immunoblotting, and immunocytochemistry.

| Experiment | Antibody (Conjugate) | Source | Dilution |
| --- | --- | --- | --- |
| IP; Immunoblotting | Mouse α-Flag | Sigma-Aldrich #1804 | 1:250 (IP) |
| | Rabbit α-DYKDDDDK | Cell Signaling #14793 | 1:750 |
| | Rabbit α-myc | Cell Signaling #2278 | 1:750 |
| | Goat α-mouse (HRP) | Promega #W402B | 1:2500 |
| | Goat α-rabbit (HRP) | Promega #W401B | 1:2500 |
| Immunocytochemistry | Mouse α-Flag | Sigma-Aldrich #1804 | 1:300 |
| | Rabbit α-At1a | Abcam #76020 | 1:500 |
| | Rabbit α-CNX | Abcam #92573 | 1:1000 |
| | Goat α-rabbit (Alexa Fluor 594) | Thermo Fisher #A11001 | 1:1000 |
| | Goat α-mouse (Alexa Fluor 488) | Thermo Fisher #A11012 | 1:1000 |

### Immunoblotting

Input lysates (5 μg of total protein) and entire elution products were combined with 4x Laemmli buffer (Bio-Rad) supplemented with 10% β-mercaptoethanol (Sigma-Aldrich, St Louis, MO). Samples were subsequently incubated at 37°C for 20 minutes to avoid boiling-induced aggregation of multi-pass transmembrane proteins (16). Samples were separated using 4-15% gradient SDS-PAGE gels (Bio-Rad), transferred to PVDF membranes (Bio-Rad), and blocked in 3% skim milk powder (BioShop) for one hour at room temperature with agitation. Blots were incubated with primary antibodies at the appropriate concentrations in blocking solution overnight at 4°C with agitation. The next day, blots were washed three times with wash buffer (TBS supplemented with 0.2% Tween-20) and incubated with horseradish peroxidase-conjugated secondary antibodies at the appropriate concentrations in blocking solution for one hour at room temperature with agitation. Blots were then washed three additional times with wash buffer, followed by chemiluminescence detection (Thermo Fisher Scientific) using the ChemiDoc imaging system (Bio-Rad). Antibody concentrations are provided in Table 4.

### Immunocytochemistry

Sterile 13 mm coverslips (Sarstedt) were inserted into 24-well culture plates and coated with 100 μg/mL poly-D-lysine solution (Sigma-Aldrich) according to the manufacturer’s instructions. HEK293T cells were re-seeded at low dilutions onto coated coverslips. The next day, Lipofectamine 3000 was utilized according to the manufacturer’s instructions to co-transfect 10 ng of *3xFlag-Ptchd1* expression plasmids and 490 ng of the inert carrier plasmid pBV-Luc (www.addgene.org #16539). 24 hours later, cells were rinsed with PBS and fixed with ice-cold 100% methanol at −20°C for 15 minutes. Cells were then washed twice with PBS and incubated with blocking solution [10% goat serum (Cell Signaling) in PBS supplemented with 0.1% Tween-20] for one hour at room temperature with agitation. After blocking, cells were incubated with primary antibodies at the appropriate concentrations in blocking solution overnight in a humidified chamber at 4°C with agitation. The next day, cells were washed three times with PBS, and incubated with Alexa Fluor-conjugated secondary antibodies at the appropriate concentrations in blocking solution for one hour at room temperature with agitation in the dark. Cells were then washed three additional times with wash buffer, followed by incubation with NucBlue reagent (Thermo Fisher Scientific) according to the manufacturer’s instructions. Finally, cells were mounted on glass slides using Dako mounting medium (Agilent). Antibody concentrations are provided in Table 4. Three biological replicates were performed for immunocytochemistry experiments. For transfected N2a cells, cells grown on coverslips were fixed in 4% PFA at room temperature for 10 minutes, permeabilized with 0.1% TritonX-100 in PBS, followed by blocking with 0.5% BSA in PBS for 30 minutes at RT. The cells were incubated with primary antibodies in 0.5% BSA in PBS solution for 1hr at room temperature or overnight at 4°C. After washing 3 times with 0.1% TritonX-100 in PBS, the cells were then incubated with AlexaFluor secondary antibodies for 45 minutes at room temperature. The cells were then washed washing 3 times with 0.1% TritonX-100 in PBS and then mounted onto slides using VectaShield Mounting media with DAPI (Vector Labs, #64335). Images were taken using the Nikon Eclipse 80*i* fluorescence microscope using QICAM-UV Fast 1394 camera (QImaging, BC, Canada) and QCapture Suite PLUS software.

### Quantification of co-localisation PtchD1 in HEK293 cells

Images were acquired using a Leica TCS SP8 confocal microscope and the Leica Application Suite X software. The 405 nm (66.6% intensity), 488 nm (5.2% intensity), and 552 nm (2.0% intensity) lasers were used. Images were acquired sequentially under 63x magnification. For each image, consecutive high-resolution z-stacks were acquired with a z-interval of 0.6 μm. For each variant, three biological replicates (each consisting of three consecutive z-stacks for 6-7 separate images) were blindly analyzed by an independent technician. Pixel intensity thresholds were adjusted identically for all images: 1) 3xFlag-Ptchd1 (minimum 30; maximum 255) and 2) ER marker Calnexin (CNX) or cell membrane marker ATP1A1 (minimum 15; maximum 230). To quantify the overlapping intensity values of 3xFlag-Ptchd1 and CNX or ATP1A1, the Pearson correlation coefficient (PCC) was computed with the JACoP (17) plugin in FIJI using default settings. To calculate statistical significance, a one-way analysis of variance (ANOVA) was used, followed by a Tukey’s Honestly Significant Difference (HSD) test to compare variants of Ptchd1 with wildtype.

## Results

### Identification of SNAPIN as a PTCHD1-Binding Protein

In order to identify novel neural proteins that may be putatively interacting with predicted exoplasmic domains of PTCHD1, lumenal loop 1 or a fusion of lumenal loops 1 and 2 were separately used as bait in yeast two-hybrid screens. Subsequent sequencing of positive clones from the screen of lumenal loop 1 against an adult human brain cDNA library indicated highly successful bait-prey interaction between this region of PTCHD1 and SNAPIN, with 17 hits, compared to the next highest candidate protein, with two hits (Table S1). This result was confirmed via co-IP, which revealed that mouse Snapin interacts strongly with the mouse Ptchd1 lumenal loop 1, and very weakly with the lumenal loop 1-loop 2 fusion protein (Figure 1A). To exclude the possibility that the binding between human PTCHD1 and human SNAPIN is an artificial byproduct of their spontaneous proximity to one another during cell lysis or ectopic expression in yeast cells, immunocytochemistry was used to evaluate their subcellular localization in P19-induced neural cells. Both PTCHD1 and SNAPIN were observed to co-localize within dendritic projections (Figure 1B).

**Figure 1.**
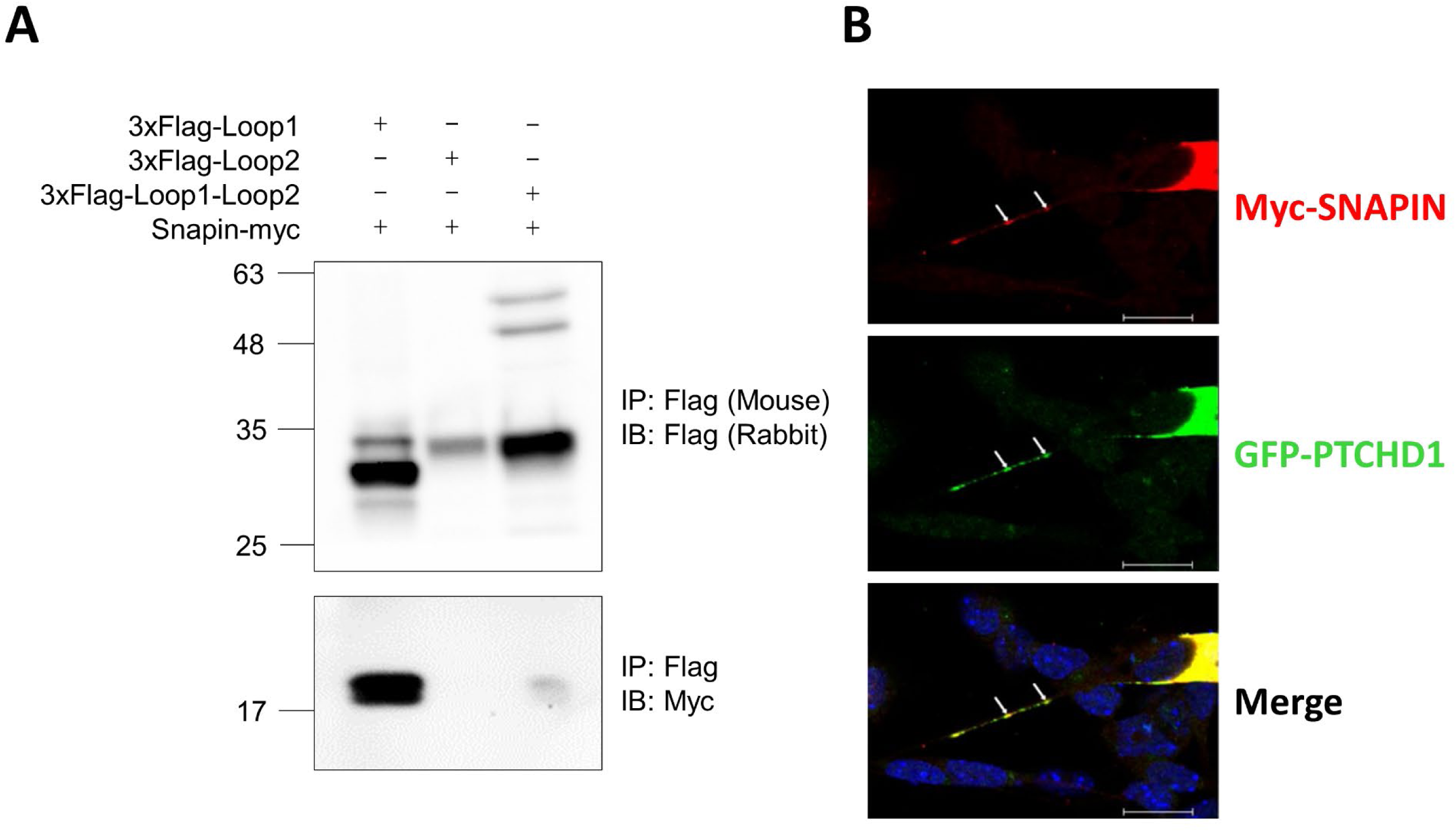
Protein interaction and neuronal co-localization between PTCHD1 and SNAPIN. **A)** Immunoblot image following co-IP of 3xFlag-tagged mouse Ptchd1 lumenal loop 1, lumenal loop 2, or a lumenal loop1-loop 2 chimera (*top blot*); and myc-tagged mouse Snapin (*bottom blot*). **B)** Immunocytochemical staining of transiently expressed GFP-tagged human PTCHD1 and myc-tagged human SNAPIN in P19-induced neurons after six days of differentiation. Arrows indicate a region along the dendritic process where PTCHD1 and SNAPIN exhibit co-localization. Scale bar represents 20 μm.

### Binding of Snapin to Clinical Variants

*PTCHD1* single nucleotide variants evaluated in this study were identified in the literature (6,8,18,19), in clinical genomics databases (11,20), and through personal communication with clinicians (Table 2). Two of these variants, p.Gln102Arg and p.Val150Met, were identified in multiplex ID Pakistani families (Figures S3 and S4, respectively). In order to ascertain if the mechanism of pathogenicity for clinically-identified point mutations within *PTCHD1* is related to an impaired capacity to bind SNAPIN, co-IP experiments were performed with these variants using tagged mouse *Ptchd1* and *Snapin* expression constructs in HEK293T cells. It was observed that all lumenal loop variants evaluated in this study were both detectable at the protein level and able to interact with Snapin (Figure 2). Variations in monomeric protein expression were observed, with the variants p.Pro75Leu, p.Pro75Gln, p.Val195Ile, p.Tyr213Cys, and p.Phe549Cys exhibiting attenuated protein levels at the expected size relative to wildtype Ptchd1 (Figure 2). The two transmembrane domain (TMD) variants, P32R and G303R, were not tested experimentally here, as previous studies had shown reduced expression levels for constructs with these mutations (10, 16).

**Figure 2.**
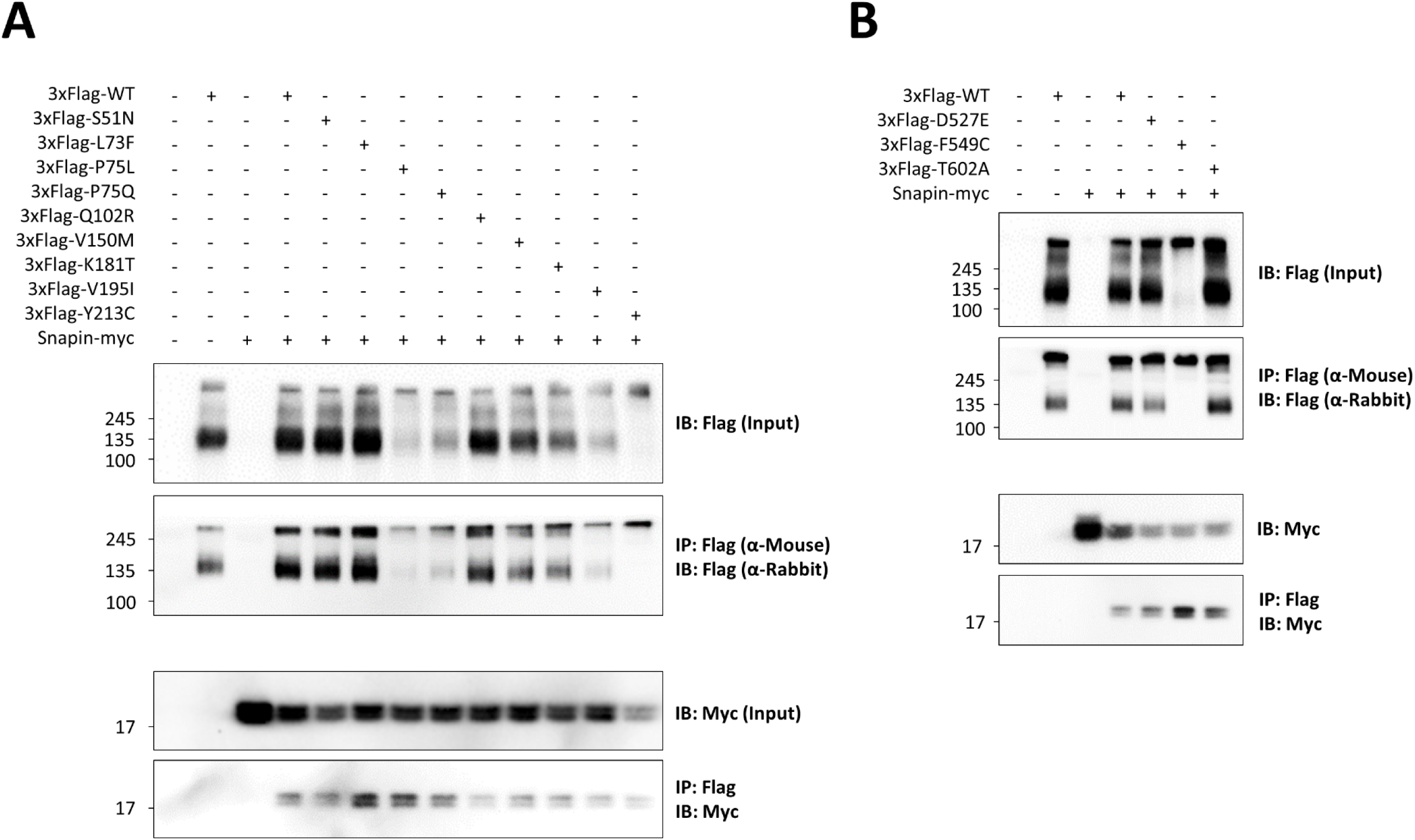
Protein interaction of clinical lumenal loop missense variants and Snapin. Immunoblot images following co-IP of 3xFlag-tagged Ptchd1 **A)** lumenal loop 1 and **B)** lumenal loop 2 missense variants and myc-tagged Snapin. Input blots are shown above their respective Flag-immunoprecipitated blots.

### Neuronal and Non-Neuronal Subcellular Localization of Clinical Variants

As the PTCHD1 VUS missense variants do not appear to hinder binding of PTCHD1 to SNAPIN, efforts to understand their relationship to pathogenicity next focused on evaluation of their subcellular localization. Upon transient transfection in HEK293T cells, six of the 12 the lumenal point mutants p.Pro75Leu, p.Pro75Gln, p.Val195Ile, p.Tyr213Cys, p.Asp527Glu, and p.Phe549Cys were all observed to demonstrate increased retention within the ER in comparison with wildtype (Figure 3). Mutations at p.Pro75 appear to be particularly deleterious, with p.Pro75Leu and p.Pro75Gln exhibiting 75% and 84% higher associations with the ER marker Calnexin (CNX), respectively, than wildtype (Figure 3B). Correspondingly, all of these variants, with the exception of p.Asp527Glu, were associated with attenuated localization within the cell membrane relative to wildtype (Figure 4). Variants generating cysteine residues, undesirable because of the potential to generate additional disulfide bridging, thus impacting 3D structure, seemingly impaired cell membrane trafficking to a significant extent, with p.Tyr213Cys and p.Phe549Cys displaying 67% and 68% less co-localization with the cell membrane marker ATP1A1, respectively, than wildtype (Figure 4B). Similarly, mutations affecting p.Pro75 (proline residues are particularly rigid within a 3D structure) considerably impaired the ability of these point mutants to be trafficked to the cell membrane, as co-localization with ATP1A1 was attenuated in p.Pro75Leu and p.Pro75Gln by 43% and 42%, respectively, relative to wildtype (Figure 4B).

**Figure 3.**
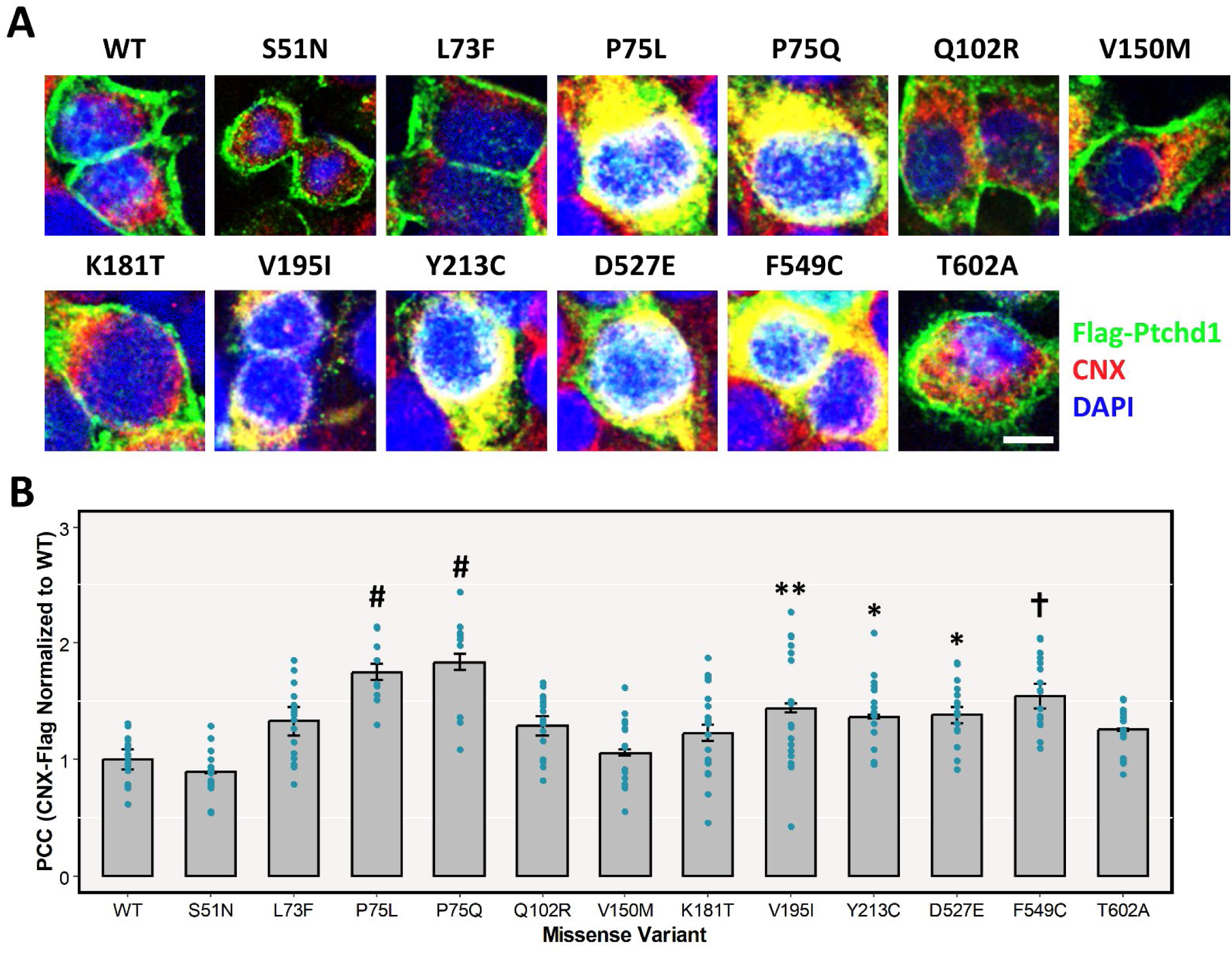
ER retention of clinical missense variants. **A)** Immunocytochemical staining of exogenous Flag-Ptchd1 (green), endogenous CNX (ER marker; red signal), and DAPI (nuclear marker; blue signal) in HEK293T cells. Representative merged fluorescent images from three independent experiments were taken for wildtype and each missense variant. **B)** PCC, calculated by JACoP, of overlapping pixel intensities between transiently expressed 3xFlag-tagged Ptchd1 and endogenous CNX (ER marker). Data are expressed as the mean ± SEM, and each missense variant was normalized to wildtype Ptchd1. Individual data points for all technical replicates are plotted. Data were analyzed using a one-way ANOVA followed by a Tukey’s HSD test (* p < 0.05; ** p < 0.01; † p < 0.001; # p < 0.0001; n = 3 immunocytochemical co-staining experiments from three separate transfections with 6-7 images captured per staining; d.f. = 2). Scale bar represents 10 μm.

**Figure 4.**
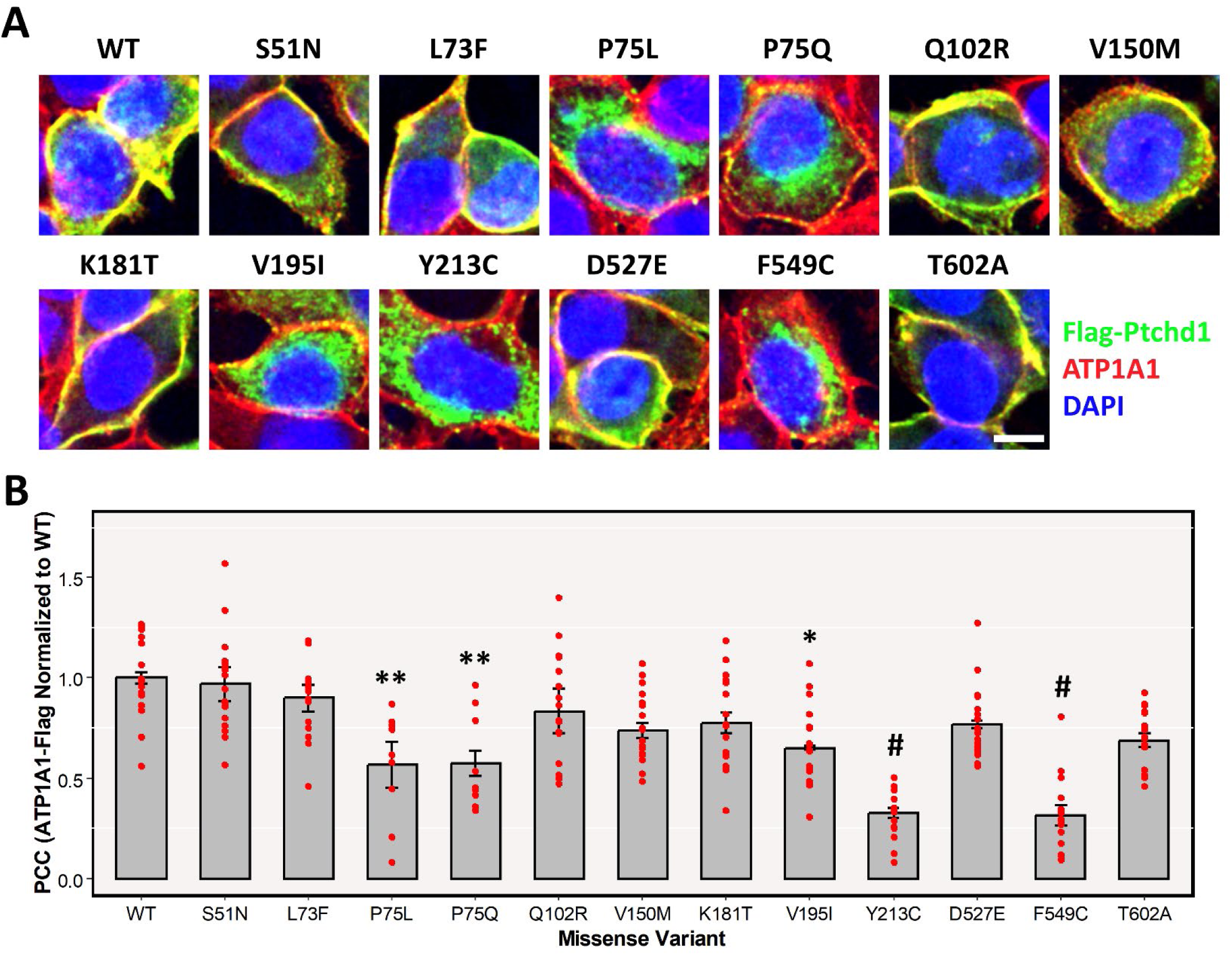
Membrane trafficking of clinical missense variants. **A)** Immunocytochemical staining of exogenous Flag-Ptchd1 (green signal), endogenous ATP1A1 (membrane marker; red signal), and DAPI (blue signal) in HEK293T cells. Representative merged fluorescent images from three independent experiments were taken for wildtype and each missense variant. **B)** PCC, calculated by JACoP, of overlapping pixel intensities between transiently expressed 3xFlag-tagged Ptchd1, and endogenous ATP1A1 (membrane marker), in HEK293T cells. Data are expressed as the mean ± SEM, and each missense variant was normalized to wildtype Ptchd1. Individual data points for all technical replicates are plotted. Data were analyzed using a one-way ANOVA followed by a Tukey’s HSD test (* p < 0.05; ** p < 0.01; # p < 0.0001; n = 3 immunocytochemical co-staining experiments from three separate transfections with 6-7 images captured per staining; d.f. = 2). Scale bar represents 10 μm.

We next sought to characterize trafficking of lumenal loop1 and 2 variants (plus P32R and G303R from TMD1 and 3 respectively) in a neuronal system. Transfections of these variants into Neuro-2A (N2A) cells revealed categorical patterns of localisation (Figure 5). Transient expression of wild type Ptchd1 results localization to neuronal processes of N2A cells. In these processes, Ptchd1 appears in a punctate pattern along the processes as well as in the small spikes that emanate perpendicular to these processes (**Figure 5A**). While Snapin expression is also observed in these same processes, Snapin is confined to the middle of the processes and specific co-localisation with punctate pattern of wild type Ptchd1 is not evident. The localisation of various Ptchd1 variants was then assessed. Two specific patterns of expression were observed for the variants. The variants p.Ser51Asn (Figure 5B; upper panels), p.Gln102Arg, p.Val195Ile, and p.Asp527Glu trafficked to plasma membranes of both the cell body and dendrites in a punctate pattern indistinguishable from wildtype Ptchd1. A distinct category of localisation was seen for other variants, as illustrated by the p.Pro32Arg variant (Figure 5B; lower panels). These variants were confined to the cell body and showed no evidence of trafficking to the processes when transiently expressed. Variants behaving in this manner included the lumenal variants p.Pro75Gln, p.Val150Met, p.Tyr213Cys, and p.Phe549Cys, as well as the transmembrane variants (Table 5).

**Figure 5.**
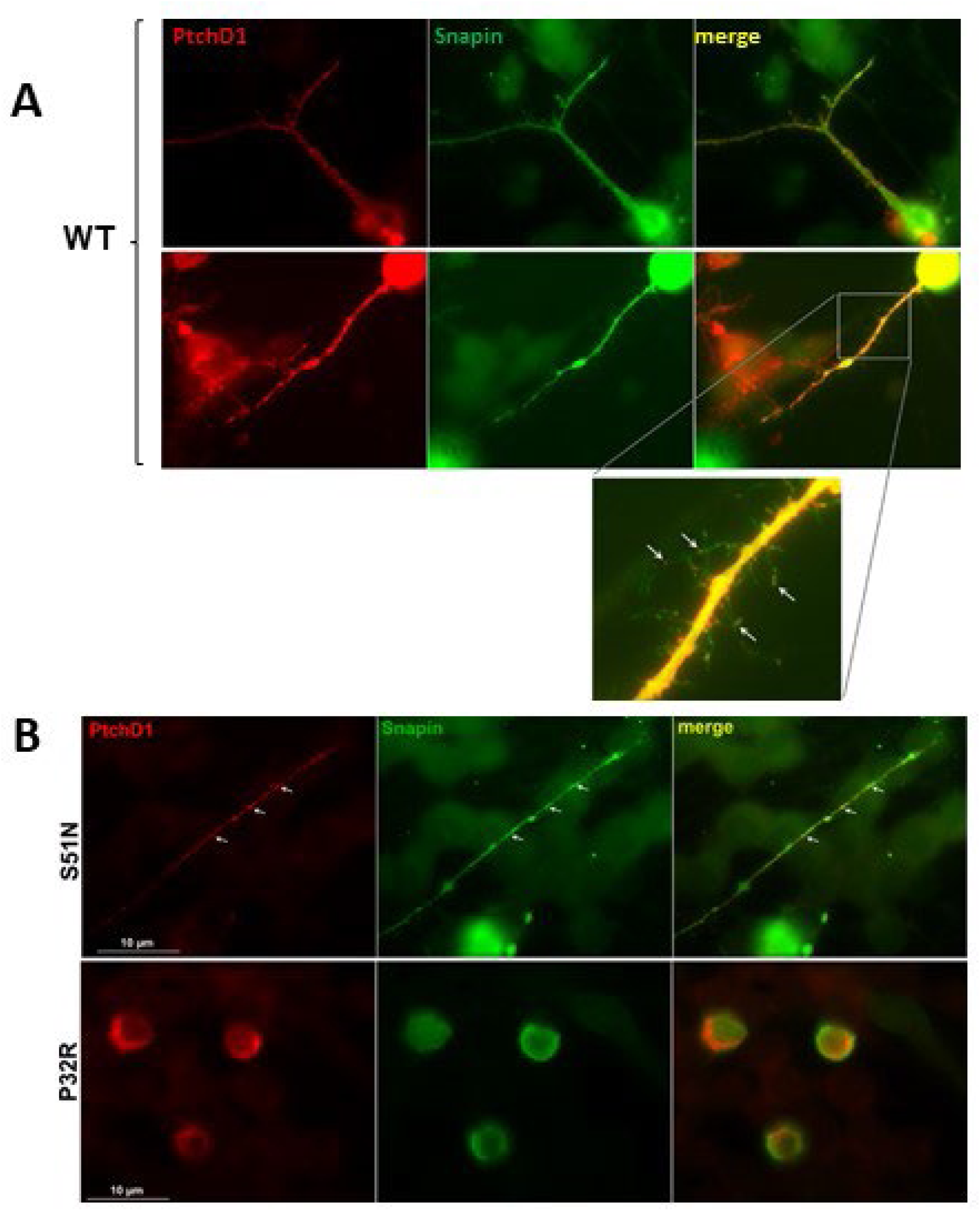
Localisation of Ptchd1 and Snapin in Neuro2A cells. Wild type Ptchd1 **(A)** or Ptchd1 missense variants **(B)** were co-expressed with Snapin in N2a cells. **(A)** Wild type Ptchd1 localizes to neuronal processes appearing in a punctate pattern along processes. The enlarged inset shows Ptchd1 localizes in discrete structures in neuronal spikes along the length of these processes. Snapin also localizes to the same structures but in a diffuse pattern that, while overlapping, does not appear as specific puncta. **(B)** Variants of PtchD1 have distinct localization qualities. A subset of variants, such as S51N (upper panels), localize in a manner indistinguishable from wild type PtchD1. A distinct subset of PtchD1 variants, including P32R (lower panels), were found localized only to the cell body of N2a cells and not in neuronal processes (see Table 5, summarizing all missense variant data).

**Table 5.** Summary of subcellular trafficking of missense variants in neuronal cells. Here, for comparison purposes, transmembrane domain (TMD) missense variants P32R and G303R were also used.

| **Mutation** | **Location** | **Membrane**<br>**Localization** | **Dendrite**<br>**Localization** |
| --- | --- | --- | --- |
| Wildtype |  | + | Punctate pattern |
| P32R | TMD 1 | - | - |
| S51N | Loop 1 | + | Punctate pattern |
| P75Q |  | - | + |
| Q102R |  | + | Punctate pattern |
| V150M |  | - | - |
| V195I |  | + | Punctate pattern |
| Y213C |  | - | + |
| G303R | TMD 3 | - | - |
| D527E | Loop 2 | + | Punctate pattern |
| F549C |  | - | - |

Taken together, while we have shown the association of Ptchd1 with the important adaptor of SNARE complexes, Snapin, missense variants of Ptchd1derived from ASD/ID patients exhibit distinct patterns of subcellular localisation. In the case of neurons, a number of these variants fail to localize to the neuronal processes where it is expected that Ptchd1 activity is important thus effectively causing a functional “knock-out” of Ptchd1 in these cells.

## Discussion

Due to the paucity of definitive functional information for PTCHD1, studies have sought to deduce its role by delineating its network of interacting proteins. Affinity purification experiments with adult mouse neural lysates indicate that Ptchd1 binds with the PSD proteins Sap102, Psd95, Dlg1-3, Magi1, Magi3, and Lin7, as well as with components of the retromer complex Snx27, Vps35, and Vps26b (7,10). Of these, the only interactions with Ptchd1 that have been reported to occur independent of the C-terminal PDZ-binding domain is with the retromer complex proteins Vps35 and Vps26b (10). Given the predicted multi-pass transmembrane structure and membrane topology of PTCHD1, it is plausible that in addition to the cytoplasmic C-terminus, the two large lumenal loops may facilitate further protein-protein interactions. In this regard, the present study has identified a novel binding partner, SNAPIN, which demonstrates a high-affinity interaction with the first lumenal loop of PTCHD1.

SNAPIN associates with Synaptosomal-associated protein 25 (SNAP25) (21), which is a core component of the *trans*-SNARE complex, along with Synaptobrevin and Syntaxin (22). The *trans*-SNARE complex facilitates the fusion of exocytic vesicles, as well as the exocytosis of neurotransmitter-containing synaptic vesicles following depolarization of pre-synaptic neurons (23). In *C. elegans*, binding of the SNAPIN ortholog SNPN-1 to SNAP25 stabilizes SNARE complex formation and promotes synaptic vesicle priming (24). Functionally, primary cortical neurons from Snapin^null^ mice show reduced frequency of mEPSCs, a smaller pool of release-ready synaptic vesicles, and de-synchronized synaptic vesicle fusion (25). In addition, at the neuromuscular junctions of SNPN-1^null^ *C. elegans*, the amount of docked, fusion-competent vesicles was significantly reduced, although the overall kinetics of synaptic transmission were unaffected (24). Finally, the dosage of Snapin in mouse primary hippocampal neurons appears to influence dendritic arborization by its binding to the cytosolic PSD95 interactor Cypin [mouse ortholog of human guanine deaminase (GDA)] (26). Collectively, these data indicate that SNAPIN is intimately involved in both neurodevelopment and neurotransmission. Developmentally, Snapin^null^ mutant mice exhibit embryonic lethality (24), whereas Ptchd1 mutant mice are viable, but demonstrate an array of neurodevelopmental abnormalities and ASD-like behaviours (5). This disparity indicates that in the absence of Ptchd1, Snapin still maintains a sufficient basal level of activity to ensure embryonic survival. Studies have indicated that dysregulation of neurotransmission is implicated in the pathoetiology of ASD (27). Therefore, PTCHD1 may be involved in coordinating SNAPIN-mediated synaptic vesicle exocytosis, contributing to the precise synchrony of neurotransmission within the brain that is required for proper cognition.

Identification of SNAPIN as a putative PTCHD1 binding partner was obtained from a yeast two-hybrid screen. An inherent limitation with this *in vitro* approach is that it provides artificial opportunities for protein interaction, potentially resulting in false positives as the candidate proteins are expressed ectopically within the same subcellular compartment. Snapin has been found to be expressed ubiquitously in various rat brain sub-regions, including both the cerebellum and cortex (21), where *PTCHD1* is abundant at the transcript level (5). Furthermore, in fractionated rat cerebral synaptosomes, Snapin was exclusively detected in synaptic vesicles (21). Together, these data are consistent with the subcellular co-localization observed between PTCHD1 and SNAPIN within the dendritic processes of P19-induced neurons in this study (Figure 1B). In addition to this neuronal co-localization, the validated binding between Snapin and the first lumenal loop of Ptchd1 (Figure 1A) suggests the likelihood of a genuine functional interaction.

This study focused primarily on nonsynonymous variants within the soluble lumenal loops of PTCHD1. Ten of the lumenal missense variants were detectable in monomeric form at varying levels (Figure 2). This is in contrast to Halewa et al., where marked instability of the p.Lys181Thr point mutant was reported (8), a result that was not observed in this study (Figure 2). This discrepancy may be attributable to differences in the epitope tag, transfection conditions, or the length of the post-transfection incubation period prior to cell lysis. Furthermore, the two missense variants that generated non-canonical cysteine residues, p.Tyr213Cys and p.Phe549Cys, were only detectable in aggregate form (Figure 2), possibly due to the formation of disulfide bridge cross-linked homodimers. Finally, the two variants involving mutated proline residues, p.Pro75Leu and p.Pro75Gln, as well as the variant p.Val195Ile, appeared to demonstrate reduced monomeric protein expression (Figure 2). Within a protein, the conformational rigidity of proline residues confers unique local secondary structure, with the cyclic nature of proline side chains anchoring the dihedral angle φ at approximately −65° (28). Consequently, substitutions generating or abolishing proline residues may result in relatively pronounced local conformational changes in comparison with other types of substitutions (29), leading to protein misfolding and aggregation.

Immunocytochemical analyses revealed that six missense variants (p.Pro75Leu, p.Pro75Gln, p.Val195Ile, p.Tyr213Cys, p.Asp527Glu, and p.Phe549Cys) displayed apparent retention by the ER (Figure 3). Consistent with this, five of these ER-retained missense variants also exhibited decreased plasma membrane localization, with the only exception being p.Asp527Glu (Figure 4). The protein trafficking data for p.Phe549Cys are in concordance with findings from Xie et al., who reported that this variant demonstrated an inability to be fully N-linked glycosylated (14), which would therefore lead to apparent retention by the ER. However, these authors also observed complete N-linked glycosylation of p.Pro75Gln (14), which is incongruent with the ER retention that was evident for that missense variant in this study. An explanation for this disparity may arise from their fusion of GFP to the N-terminus of PTCHD1, which, given its large size relative to the 3xFlag epitope tag used in this study, may have an intrinsic stabilizing effect on certain missense variants. Functionally, the interaction between Snapin and the first lumenal loop of Ptchd1 was not abolished by any of the nine clinical point mutations evaluated in this study (Figure 2). This implies that the Snapin-binding domain of Ptchd1 is not structurally reliant on these specific amino acid residues. Clinically, the findings from this study suggest that the pathoetiologies of the missense variants evaluated in this study are unrelated to their interactions with SNAPIN.

An inherent constraint with the use of HEK293T cells as a model for protein expression in the context of ASD and ID is that it is non-neuronal, which may affect the neuropathological translatability of these findings. This concern is minimized by the fact that mechanisms of retention of misfolded proteins by the rough ER are highly conserved across cell types, and that this system has previously been used to evaluate the subcellular localization of PTCHD1 missense variants

(8). To overcome this limitation, we also sought to evaluate PTCHD1 subcellular trafficking in a neuronal context in this study. In neurons derived from differentiated N2A cells, wildtype PTCHD1 is trafficked to plasma membranes of both the cell body and dendrites, and also displays a punctate localization pattern within dendrites (Figure 5). Of the subset of missense variants similarly assayed in this study, both p.Ser51Asn and p.Gln102Arg were appropriately trafficked to the plasma membrane of differentiated N2A cells, while p.Pro32Arg, p.Pro75Gln, p.Val150Met, p.Tyr213Cys, p.Gly303Arg, and p.Phe549Cys all qualitatively showed an inability to localize to the plasma membrane (Figure 5; Table 5). With the exception of p.Val150Met, these data are consistent with our observations from HEK293T cells (Figure 4), or as reported by our group previously (14). Interestingly, p.Val195Ile demonstrated plasma membrane localization in our neuronal system, but exhibited ER retention and diminished membrane trafficking in HEK293T cells (Figures 3 and 4). This discrepancy may suggest that neuronal cells possess a unique mechanism to allow some point mutants, despite relative instability, to exit the ER and be trafficked to their proper cellular destination.

The subcellular localization data from the present study offer a pathoetiological mechanism for a number of these missense variants, in which protein misfolding-mediated aggregation and retention in the rough ER, and subsequent impaired dendritic membrane trafficking, leads to insufficient bioavailable PTCHD1 at the PSD. Indeed, chronic and excessive ER stress has also been suggested to be associated with ASD pathology, potentially through impaired membrane trafficking of GABA receptor subunits (30) and constitutive activation of mammalian target of rapamycin (mTOR) mediated by tuberous sclerosis complex loss of function (31). The mechanism of pathogenicity for the other missense variants evaluated in this study that displayed canonical subcellular localization (p.Ser51Asn, p.Leu73Phe, p.Gln102Arg, p.Val150Met, p.Lys181Thr, and p.Thr602Ala) remains elusive. Loop regions of transmembrane proteins constitute binding pockets for interacting proteins and other biomarkers (32), and therefore it is feasible that the binding of either cholesterol or a still-unidentified PTCHD1 interacting partner may be affected by these point mutations. Collectively, these findings provide a platform for future diagnostic assays and improved interpretation of PTCHD1 VUSs by genetic diagnosticians.

This study has identified and validated a novel PTCHD1 interacting partner, SNAPIN, and confirmed that selected clinical missense variants within the lumenal loops do not abolish this interaction. In addition, we report that several missense variants exhibit ER retention, as well as display impaired plasma membrane localization in neurons. Future research should be aimed at clarifying the effects of PTCHD1 on SNAPIN-mediated synaptic vesicle exocytosis; further elucidating the endogenous network of proteins interacting with PTCHD1, possibly through biotin identification; and quantifying the stability of PTCHD1 missense variants using highly sensitive biochemical techniques, such as circular dichroism, differential scanning calorimetry, or microscale thermophoresis.

## Acknowledgements

This work was supported by a grant from the Canadian Institutes of Health Research to J.B.V. (#PJT-156367).

