## Supplementary Materials for "Autism-associated PTCHD1 missense variants bind to the SNARE-associated protein, SNAPIN, but exhibit impaired subcellular trafficking"

**Supplementary Material**

**Contents:**

**Figure S1 and S2: Sanger sequence validation of PTCHD1 missense constructs generated using site-directed mutagenesis.**

**Figure S3: Pedigree and Sanger validation of Q102R variant in multiplex family from Pakistan.**

**Figure S4: Pedigree and Sanger validation ofV150M variant in multiplex family from Pakistan.**

**Figure S5: Clustal 2.1 alignment of PTCHD1 across vertebrate evolution, showing conservation at missense variants.**

**Table S1: Full results from PTCHD1 yeast two-hybrid screen.**

**Table S2: Allele frequencies and clinical predictions of PTCHD1 missense variants.**

**Table S3: *In silico* predictions for pathogenicty of PTCHD1 missense variants.**

**Supplementary References**

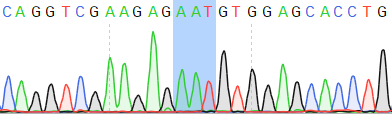

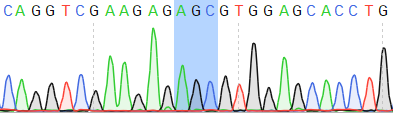

**WT**

**S51N**

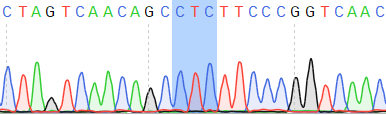

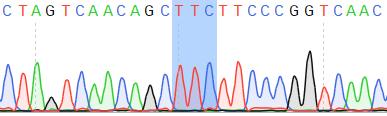

**WT**

**L73F**

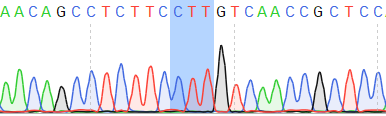

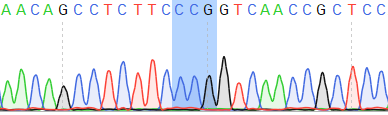

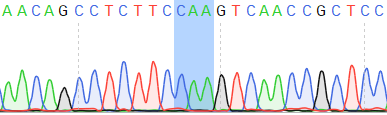

**WT**

**P75L**

**P75Q**

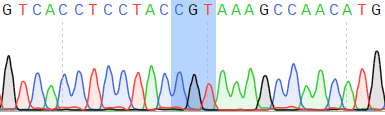

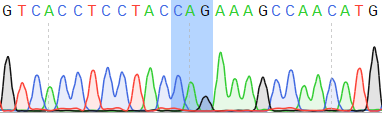

**WT**

**Q102R**

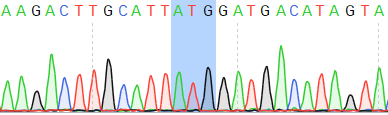

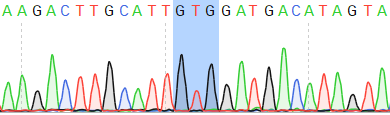

**WT**

**V150M**

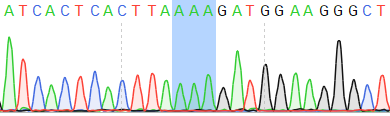

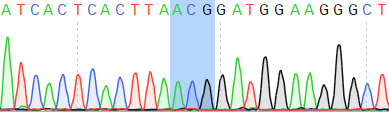

**WT**

**K181T**

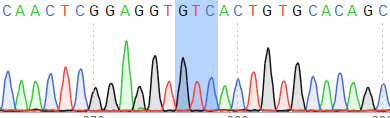

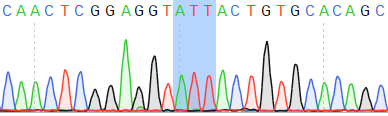

**WT**

**V195I**

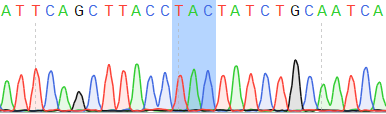

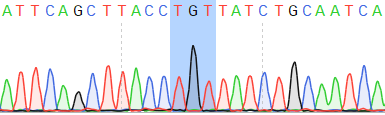

**WT**

**Y213C**

**Figure S1. DNA electropherograms of lumenal loop 1 missense variants.**

Mutant codons are highlighted in blue. Missense variants p.Ser51Asn, p.Leu73Phe, p.Pro75Leu, p.Pro75Gln, p.Gln102Arg, p.Val150Met, p.Lys181Thr, p.Val195Ile, and p.Tyr213Cys are located in lumenal loop 1

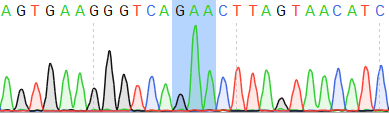

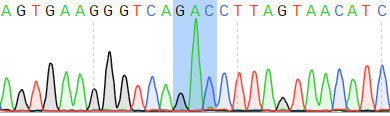

**WT**

**D527E**

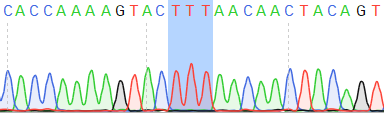

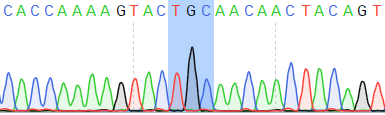

**WT**

**F549C**

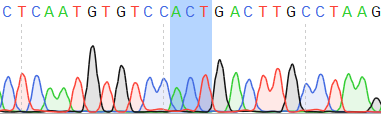

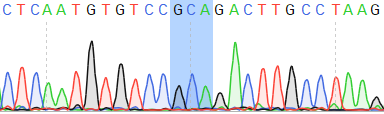

**WT**

**T602A**

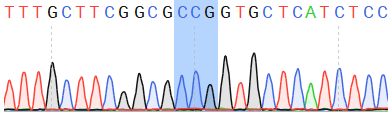

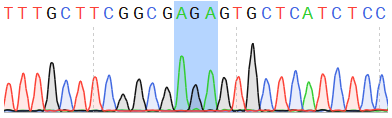

**WT**

**P32R**

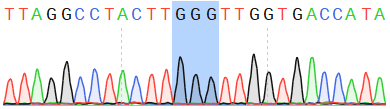

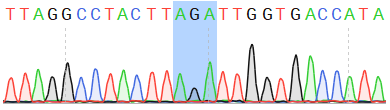

**WT**

**G303R**

**Figure S2. DNA electropherograms of lumenal loop 2 and TMD missense variants.**

Mutant codons are highlighted in blue. Missense variants p.Pro32Arg and p.Gly303Arg are located in the first and third TMDs, respectively; missense variants p.Asp527Glu, p.Phe549Cys, and p.Thr602Ala are located in lumenal loop 2

**A/G**

**A**

I:1

I:2

II:1

II:2

II:3

II:4

II:5

**A/A**

**G**

**G**

**n.a.**

**n.a.**

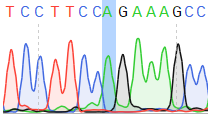

**c.305A**

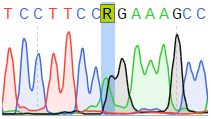

**Heterozygous**

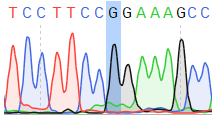

**c.305G**

**A**

**B**

**Figure S3. SNV Identification (c.305A > G; p.Gln102Arg) in a multiplex ID Pakistani family.**

**A)** Pedigree of Family ARSID_M4 with two males and one female diagnosed with ID. Both affected males (II:2 and II:3) contain a point mutation (c.305A > G; p.Gln102Arg), which was inherited from their unaffected mother (I:1). **B)** Electropherograms showing the canonical *PTCHD1* sequence *(top)*, the c.305A > G point mutation *(bottom)*, and the heterozygous maternal carrier *(middle)*.

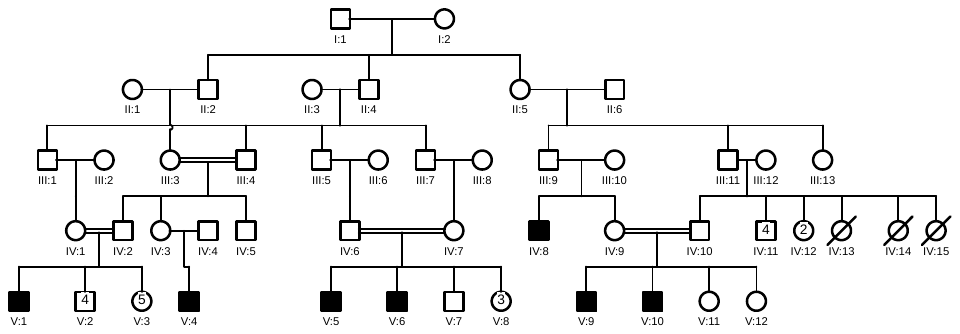

**(G)**

**(G)**

**G/G**

**G**

**A**

**G/G**

**G**

**(G/A)**

**G**

**G**

**G**

**G**

**G**

**G**

**G**

**G**

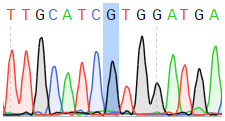

**c.448G**

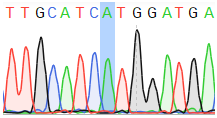

**c.448A**

**A**

**B**

**Figure S4. SNV Identification (c.448G > A; p.Val150Met) in a multiplex ID Pakistani family.**

**A)** Pedigree of Family AS30 with seven males diagnosed with ID. One of these affected males (IV:8) contains a point mutation (c.448G > A; p.Val150Met), which is *de novo* or was inherited from his unaffected mother (III:10). **B)** Electropherograms showing the canonical *PTCHD1* sequence *(top)* and the c.448G > A point mutation *(bottom)*.

**Figure S5. Evolutionary sequence alignments for PTCHD1 using CLUSTAL 2.1.** Protein sequences from representative species spanning >400 million years of the vertebrate lineage, including: human (NP_775766.2), mouse (NP_001087219.1), Tasmanian devil (XP_003763547.1), alligator (XP_006266959.1), Xenopus (XP_004911719.1), zebrafish (XP_690754.1), coelacanth (XP_006013422.1), and lamprey (Ensembl gene prediction ENSPMAT00000029719.1). Residues corresponding to missense variants evaluated in this study are indicated with red highlight.

```
								  P32R	 	     S51N
human           MLRQVLHRGLRTCFSRLGHFIASHPVFFASAPVLISILLGASFSRYQVEESVEHLLAPQH
mouse           MLRQVLHRGLRTCFSRLGHFIASHPVFFASAPVLISILLGASFSRYQVEESVEHLLAPQH
Tas_devil       MLRQVLHRGLRTSFSRLGHFIASHPVFFASAPVLISILLGASFSRYQIEESVEHLLAPKH
alligator       MLRQVLHRGLRTSFSRLGHFVASHPVFFASAPVLISILLGASFSRYHIEESVESLLAPKH
canary          MLRQVLHRGLGTSFSRLGHFVASHPVFFASAPVLISILLGASFSRYQVEESVEHLLAPTH
xenopus         MLRKVLHRGLRNCFSRLGFFIASHPVFFISAPVLISILLGASFSRYRVEENIEYLLAPKH
coelacanth      MLRQVLHKGLRTCFSRLGYFIASHPVFFASAPVLISILLGASFSRYRIEENVEYLLAPKH
zebrafish       MLRQVLHEGLRTSFHKLGHFVANHPVFFASAPVLISILLGASFSRYRIEENVEYLLAPKH
lamprey         MLGALLHAALQRCLYRLGLLVAGHPAPFLVAPALLAALLGAALSRVSVESKAEDLFAPAH
				   L73F P75Q/L			      Q102R
human           SLAKIERNLVNSLFPVNRSKHRLYSDLQTPGRYGRVIVTSFQKANMLDQHHTDLILKLHA
mouse           SLAKIERNLVNSLFPVNRSKHRLYSDLQTPGRYGRVIVTSYQKANMLDQHHTDLILKLHT
Tas_devil       SLAKIERNLVNSLFPVNRSKHRLYSDLQTPGRYGRVIITSFQQENMLDQHHTDLILKLHS
alligator       SLAKIERNLVNSLFPVNRSKHRLYSDLQTPGRYGRVIITSFRKANMLDQHHTELILKLHS
canary          SLAKIERNLVDSLFPVNRSKHRLYSDLQTPGRYGRVIITSFRKANMLDQHHTDLILKLHS
xenopus         SLAKIERNLVDSLFPVNRSKHRLYSDLQTPGRYGRVIITSSRRANMLDQHHTDLILKLHS
coelacanth      SLAKIERNLVDSLFPVNRSKHRLYSDLQTPGRYGRVIITSSRKGNMLDQFHTDLILKLHF
zebrafish       SLAKIEGNLVDSLFPVNRSKHTLYSDLQTPGRYGRVIVTS-RRGSVLDPHHVNSVLKLHN
lamprey         SLAKLEGALADELFPLERSRQRLYSELHTPGRYTRLIAVARGGGNVLDEGRRLALISLHS
						  V150M
human           AVTKIQVPRPGFNYTFAHICILNNDKTCIVDDIVHVLEELKNARATN---RTNFAITYPI
mouse           AVTKIQVPRPGFNYTFAHICVLNNDKTCIVDDIVHVLEELKNARATN---RTNFAITYPI
Tas_devil       AVTRMQVQRPGFNYTFAHICILNNDKTCIVDDIVHVLEELKTARATN---RTNFAITYPI
alligator       AVTRIQVQRPGFNYTFAHICILNNDKTCIVDDIVHVLEELKTARSSN---RTNFAITYPI
canary          AVTRIQVQRPGFNYTFAHICILNNDKTCIVDDIVHVLEELKAARSSN---RTNFAITYPI
xenopus         AVKKIQVHRPGFNYTFAHICMLSNEKTCIVDDIVHILEELKAARSQN---RTNYIITYPI
coelacanth      SVNKIQVPMLGINYTFAHMCVLNDDKTCIVDDVVHILEELQASRSLN---KTGFTIMYPI
zebrafish       TITQIQVPMLGFNYTFAHLCLLDESKSCIVDDILRVLEEMRSARASN---HSVPPLRYPI
lamprey         ALVAMPVGR---NCTFWDVCSIDWDRECVQDAIVELMGNGSSSASDASRRRSSIIIRYPK
		    K181T	  V195I		         Y213C
human           THLKDGRAVYNGHQLGGVTVHS-KDRVKSAEAIQLTYYLQSINSLNDMVAERWESSFCDT
mouse           THLKDGRAVYNGHQLGGVTVHS-KDRVKSAEAIQLTYYLQSINSLNDMVAERWESSFCDT
Tas_devil       THLKDGREVYNGHQLGGVTVHS-KDQVKSAQAVQLTYYLQTLNSLNDMVAERWESNFCDT
alligator       THLKDGREVYNGHQLGGVTVHS-KDRVKSAEAIQLTYYLQAINSLNDMVAEKWESIFCNT
canary          THLKDGREVYNGHQLGGVTVHS-KDRVKSAEAIQLTYYLQAINSLNDMVAEKWESIFCDT
xenopus         TKLKDGKEVYNGHQLGGVTVHS-KDRVKSAEAIQLTYYLQAINSLNEVVAEKWESVFCET
coelacanth      THLKNDREVYIGHQLGGVTLHS-KDRVKSAEAIQLTYYLQTINFLNDMVAEKWESIFYET
zebrafish       TKLKDGREAYIGHQLGGVLASGGRDGVRSARALQLTYYLQAVSPLNEVVAASWELLFCRE
lamprey         ARLRDNQEVYIGHQLGGVTLFQ-TDRVLSARALQITYYLDSAHAA----SELWEREFVTA

human           VRLFQK-SNSKVKMYPYTSSSLREDFQKTSRVSERYLVTSLILVVTMAILCCS-MQDCVR
mouse           VKLFQK-SNSKVKIYPYTSSSLREDFQKTSRVSERYLVTSLILVVTMAILCCS-MQDCVR
Tas_devil       VKLFQK-SNGKIKMYPYTSSTLREDFQKTSRVSERYLITSLVLMVTMAVLCCS-MQDCVR
alligator       VELFQK-SNRKVKMYPFTSSSLKEDFQKTSRVSERYLITSLVLVVSLAILCCS-MQDCVR
canary          VELFQK-SNRKVKMYPFTSSSLKEDFQKTSRVSERYLITSLVLVVTLAILCCS-MQDCVR
xenopus         VENFQK-LNREVKLYPFTSSSLGQDFQKTSRVSERYLITSLALVVSLAVICCS-MQDCVR
coelacanth      VELFQA-SNGAVKLYPFTSSSLSEDFQKTSRVSEQYLITSLILVVFLAILCCS-MRDCVR
zebrafish       LENFGK-AHPELSLHPFTSSSLQRDFQRTSRVSERPLLFSLAVCLSLAMLCCS-MRDCVR
lamprey         VERARHRHAHDLALFPFTSSSLQTDFYQSGVVAAPNLAGGCGGWWTALKNSRGRTRSEDL
			 G303R
human           SKPWLGLLGLVTISLATLTAAGIINLTGGKYNSTFLGVPFVMLGHGLYGTFEMLSSWRKT
mouse           SKPWLGLLGLVTISLATLTAAGIINLTGGKYNSTFLGVPFVMLGHGLYGTFEMLSSWRKT
Tas_devil       SKPWLGLLGLVTVSLATLTAAGIINLTGGKYNSTFLGVPFVMLGHGLYGTFEMLSSWRKT
alligator       SKPWLGLLGLLTISLATLTAAGIINLTGGKYNSTFLGIPFIMLGHGLYGTFEMLSSWRKT
canary          SKPWLGLLGLLTVTLATLTAAGIINLTGGKYNSTFLGIPFVMLGHGLYGTFEMLSSWRKT
xenopus         SKPWLGLLGLVTISLSTLTAAGIINLTGGKYNSTFLGIPFIMLGHGLYGTFEMLSSWRKT
coelacanth      SKPWLGILGLVTMSLATLTSAGIINLTGGKYNSTFLGIPFIVLGHGLYGTFEMLSSWRKT
zebrafish       TKPWLGLLALVTVSLATLTSAGILNLTGGKYNSTYLGIPFVMLGHGLFGTFEMLSSWRRT
lamprey         CFGLLVLAGFVGIALTTLATAGVMILTGTPYNSTLIIIPLVALGHGSHGATELLWTWRRL

human           RE-----DQHVKERTAAVYADSMLSFSLTTAMYLVTFGIGASPFTNIEAARIFCCNSCIA
mouse           RE-----DQHVKERTAEVYADSMLSFSLTTAMYLVTFGIGASPFTNIEAARIFCCNSCIA
Tas_devil       RE-----DQHVKERTAAVYADSMLSFSLTTAMYLVTFGIGASPFTNIEAARIFCCNSCIA
alligator       RE-----DQHVKERTAAVFADSMLSFSLTTAMYLVTFGIGASPFTNIEAARIFCCNSCIA
canary          RE-----DQHVKERTAAVFADSMLSFSLTTAMYLVTFGIGASPFTNIEAARIFCCNSCIA
xenopus         RE-----DQHVKERTAVVYADTMISFTLTTAMYLVTFGIGASPFTNIEAARIFCRNSCIA
coelacanth      RE-----DQHVKERMATVYADSMLPFSFTTAMYLVTFGIGASPFTNIEAARIFCRNSCIA
zebrafish       RE-----DQHVKERVAAVFSDCMLPFTASTALHVVTFGIGASPFTNIEAVRLFCQNACIS
lamprey         GWGARGPPAREEERLAATFARSLLPHTMLTALQVITLALGASPLTNTRAVQVFCRCACTA

human           IFFNYLYVLSFYGSSLVFTGYIENNYQHSIFCRKVPKPEALQEKPAWYRFLLTARFSEDT
mouse           ILFNYLYVLSFYGSSLVFTGYIENNYQHSIFCRKVPKPDVLQEKPAWYRFLLTARFSEET
Tas_devil       IFFNYLYVLSFYGSSLVFTGYIENNYQHSLFCRKVPKPEVLQEKPAWYRFLLTARFSEDT
alligator       IFFNYLYVLSFYGSSLVFTGYIENNYQHSIFCRKVPKPEVLQEKPAWYRFLLTAKFSEDT
canary          IFFNYLYVLSFYGSSLVFTGYIENNYQHSIFCRKVPKPEVLQEKPAWYRFLLTARFSEDT
xenopus         IFFNYLYVLSFYGSSLVFTGYIENNYQHSIFCRKVPKPEILQVKSLWYRFLMTAKFSDDT
coelacanth      VFFNYLYVLSFYGSNLVFTGYMENNYQHSLFCRKVPKPEVLQGKPLWYRLLLTAKYNEET
zebrafish       VLFNYLYILTFYGSNLVFAGYLENNYRHSLFCRRVPKPELLQQKPAWYRFLMYTHYNEEA
lamprey         TALGYCACVTFLGASSDVTPRGLQDLKTPRCICEQPDLISSKEPPCSTATTNTTNTASTA

human           AEGEEANTYES--------HLLVCFLKRYYCDWITNTYVKPFVVLFYLIYISFALMGYLQ
mouse           AEGEEANTYES--------HLLVCFLKRYYCDWITNTYVKPFVVLFYLIYISFALMGYLQ
Tas_devil       ADGEEANTYES--------HLLVCFLKRYYCDWITNTYVKPFVVLFYLIYISFALMGYLQ
alligator       TDSEETNTYES--------HLLVCFLKRYYCDWITNTYVKPFVVLFYLVYISFALMGYLQ
canary          DDSEETNTYES--------HLLVWFLKRYYCDWITNTYVKPFVVLFYLVYISFALMGYLQ
xenopus         ADAEETNSYES--------HLLVCFLKRYYCDWITNTYVKPFVILFYLVYISFALMGYLQ
coelacanth      ADSVETSTYES--------HLLICFLKRYYCDWITNTYVKPFVVLFYLVYISFALMGYLQ
zebrafish       TEAGPLCAYES--------HLLVAFMKRYYCDWITNTYVKPFVVLFYLVYVSFALMGYLQ
lamprey         AVTTTTTAAAATSGPHPPDAPPRRFMRDRYGPWISGTYVKPFVVLLYLVYASFSFMGCLQ
		     D527E			F549C
human           VSEGSDLSNIVATATQTIEYTTAQQKYFSNYSPVIGFYIYESIEYWNTSVQEDVLEYTKG
mouse           VSEGSDLSNIVATATQTIEYTTAHQKYFNNYSPVIGFYIYESIEYWNSSVQEDVLEYTKG
Tas_devil       VREGSDLSNIVATATRTIEFTTAQQKYFSNYSPVIGFYIYESIEYWNTSVQEDVLEYTKG
alligator       VNEGSDLSNIVATATRTIEYTTAQQKYFSNYSPVIGFYIYESIEYWNTSVQEDVLEYTKG
canary          VSEGSDLSNIVATATRTIEYTTAQQKYFSNYSPVIGFYIYESIEYWNTSVQEDVLEYTKG
xenopus         VHEGSDLRNIVATETRTITYTTVQQKYFSNYSPVIGFYIYESIDYWNTSVQEDVLEYTKG
coelacanth      VNEGSDLSNIVATETRTIAYTTAQQKYFSNYSPVIGFYIYESIEYWNISVQEDVLEYTKG
zebrafish       VSEGSDLSNVVATETSTIAYTRAQQRYFSSYSPVIGFYIYESIEYWNTSVQEDLLEYIKG
lamprey         LREASDLTHLVASRSATARYRAVQDRFFSDYSPVIGFYIYEPVAYWNASVQEDLLDITRR
					T602A
human           FVRISWFESYLNYLRKLNVSTGLPKKNFTDMLRNSFLKAPQFSHFQEDIIFSKKYNDEVD
mouse           FVRISWFESYLNYLRKLNVSTDLPKKNFTDMLRNSFLKTPQFSHFQEDIIFSKKYNDEVD
Tas_devil       FVRISWFESYLNYLRKLNASTGLPKKNFTDMLRNSFLKAPQFSHFSEDIIFSKKFNNEVD
alligator       FVRISWFESYLNYLRKLNISTGLPKKNFTDMLRNSFLKAPQFAHFSEDIIFSKKYNNEVD
canary          FVRISWFESYLNYLRKLNISTGLPKKNFTDMLRNSFLKTPQFAHFSEDIIFSKKYNNEVD
xenopus         FVRISWFESYLNYLRKLNMSTGLPKKNFTDILRYSFLKNPQYAHFSEDIIIPKKYNNDVE
coelacanth      FVRISWFESYLNYLRKLNMTTGLPKKNFTEMLRNSFLKTPQFSHFSEDIIFAKKYNNEVE
zebrafish       FERISWFESYLNYLHGLNITTSLSRSNFTERLRSGFLRQPRYVHFTDDIIFAKRSDGEFD
lamprey         FVTVSWLEQYTRYLRAANGTSALPRDLFVSTLCASFLRRREFAHFADDVVLAGP-PGERR

human           VVASRMFLVAKTMETNREELYDLLETLRRLSVTSKVKFIVFNPSFVYMDRYASSLGAPLH
mouse           VVASRMFLVAKTMETNREELYDLLETLRRLSVTSKVKFIVFNPSFVYMDRYASSLGAPLH
Tas_devil       VVASRMFLVAKTMETNREELYDLLETLRRLSVTSKVKFIVFNPSFVFMDRYSSSLGAPLQ
alligator       VVASRMFLVAKTMETKREELYDLLETLRKLSLTSKVKFIVFNPSFVYMDRYASSVGAPLQ
canary          VVASRMFLVAKTMETKREELYDLLETLRKLSLTSKVKFIVFNPSFVYMDRYASSVGAPLQ
xenopus         VVASRMFLVAKTMETNREELYDLLETLRKLSLTSKVKFIVFNPSFVYMDRYASSVGAPLQ
coelacanth      VVASRMFLVAKTMETKREELYDLLETLRKLSLTSKVKFIIFNPSFVYMDRYASSIGAPLQ
zebrafish       VVASRMFLIAKTTENKREEMSILLDTLRKLSLTSRVKFIIFNPSFVYMDRYASSVGAPLK
lamprey         IAASRVFMVAKTNENTREEIDGLLEALRKLSLTSRVKFTIHNPAFAFLERYALWVGAPAH

human           NSCISALFLLFFSAFLVAD--SLINVWITLTVVSVEFGVIGFMTLWKVELDCISVLCLIY
mouse           NSCISALFLLFFSAFLVAD--SLINVWITLTVVSVEFGVIGFMTLWKVELDCISVLCLIY
Tas_devil       NSCISALFLLFFSAFLVAD--SLINVWITLTVASVEFGVIGFMTLWKVELDCISMLCLIY
alligator       NSCISALFLLFFSAFLVAD--SLINVWLTLTVASVEFGVIGFMTLWKVELDCISVLCLIY
canary          NSCISALFLLFFSAFLVAD--SLINVWLTLTVASVEFGVIGFMTLWKVELDCISVLCLIY
xenopus         NSCISALFLLFFSAFLVAS--SIINVWITLTVASVEFGVIGFMTLWKVELDCISVLCLIY
coelacanth      NSCISGLFLLFFSAFLVAD--SLINVWLTVTVASVEFGVIGFMTLWKIELDCISVLCLIY
zebrafish       NSCIAALFLLFFSTFLAAD--PLVNAWLTVTVASVEFGLVGFMTLWRVELDCVSVLCLIY
lamprey         AAALAAALALLLSAGLAPPGAAPAHAWVALCAASVQFGVLGSLGLMGAQLDCAAVLCLVY

human           GINYTIDNCAPMLSTFVLGKDFTRTKWVKNALEVHGVAILQSYLCYIVGLIPLAAVPSNL
mouse           GINYTIDNCAPLLSTFVLGKDFTRTKWVKNALEVHGVAILQSYLCYIVGLFPLAAVPSNL
Tas_devil       GINHTIDNCAPLLSTFVLGKDFTRTKWVKNALETHGVAILQSYLCYIVGLIPLAAVPSNL
alligator       GINYTIDNCAPLLSTFVLGKDFTRTKWVKNALEIHGVAILQSYLCYIVGLIPLAAVPSNL
canary          GINYTIDNCAPLLSTFVLGKEFTRTKWVKNALEVHGVAILQSYLCYIVGLIPLAAVPSNL
xenopus         GINYTIDNCAPLVSTFILGKEFSRTKWVKNSLEVHGVAILQSYLCYTVGLIPLAAVPSNL
coelacanth      GINYTIDNCAPLISTFVLGKDFTRTKWVKNTLELHGVAILQSYLCYTVGLIPLAAVPSNL
zebrafish       GVNYAVDSSAPLVSAFALGRESTRTRWVKLALEQHGVPALQSYLCYGAALLPLAAVPSNL
lamprey         SLCYSAHACAPLVATFAMGRGKSRAHWTAAALDAHAAPLLHACLWFCAAVVALAAAPSNL

human           TCTLFRCLFLIAFVTFFHCFAILPVILTFLPPSKKKRKEKKNPE-NREEIECVEMVDIDS
mouse           TCTLFRCLFLIAFVTFFHCFAILPVILTFLPPSKKKRKEKKNPE-NREEIECVEMVDIDS
Tas_devil       TCTLFRCLFLIAFVTFFHCFAILPVILTFLPPSKKKRKEKKNPE-NREEIECVEMVDMDS
alligator       TRTLFRCLFLIALVTFFHCFAILPVILTFVPPSKKKRKEKKNPE-NREEIECVEMVDMDS
canary          TRTLFRCLFLIALVTFFHCFAILPVILTFLPPSKKKRKEKKNPE-NREEIECVEMVDMDS
xenopus         TRTLFRCLFLIAFVTFFHCFAILPVILTFVPPSKKKRKEKKTPE-NREEIECVEMVDLDS
coelacanth      TRTLFRCLFLIAFVTFFHCFAILPVILTFLPPSKKKKKEKKNPE-HREEIECVEMVDS--
zebrafish       TRTLFRCLFLTAIITAFHCLAILPVLLTFLPPSKKKRRERKNAAENREEIECVEMVDS--
lamprey         ARTVARCLGLASALSAVHCLVFLPVFLTICPPSKAVKRRRQAGDGEAEEGDAAGAAAGAT

human           TRVVDQITTV
mouse           TRVVDQITTV
Tas_devil       TRVVDQITTV
alligator       TRVVDQITTV
canary          TRVVDQITTV
xenopus         TRVVDQITTV
coelacanth      TRVVDQITTV
zebrafish       TRVVDQITTV
lamprey         DGAVDQATSV
```

**Table S1. Full results from PTCHD1 yeast two-hybrid screen.** Screen 1 used human PTCHD1 sequence from luminal loops 1 and 2 against an E11 mouse embryo cDNA library. Screen 2 used just PTCHD1 loop 1, against an adult human brain cDNA library.

| **Protein** | **Screen** | **Frequency of Hits** |
| --- | --- | --- |
| COX11 | PTCHD1^L1-L2^ vs. E11 Mouse Embryo cDNA Library | 3 |
| MATH1 |  | 2 |
| PAX3 |  | 1 |
| A2M | PTCHD1^L1^ vs. Adult Human Brain cDNA Library | 2 |
| ANKD49 |  | 1 |
| ANKRD55 |  | 2 |
| ATG5 |  | 1 |
| CPLX2 |  | 1 |
| DCAF17 |  | 1 |
| EPB41L1 |  | 1 |
| GFM2 |  | 1 |
| HSPD1 |  | 1 |
| KDM1A |  | 1 |
| KLHL32 |  | 1 |
| METAP2 |  | 1 |
| NOL4 |  | 1 |
| PDZD2 |  | 1 |
| PON |  | 1 |
| PSMD14 |  | 1 |
| SNAPIN |  | 17 |
| STMN2 |  | 1 |
| SYNE1 |  | 1 |
| TADA1 |  | 1 |
| TTC8 |  | 1 |
| YWHAZ |  | 1 |
| ZnF277 |  | 2 |
| ZnF350 |  | 1 |

**Table S2. Allele frequencies and clinical predictions of PTCHD1 missense variants.**

Statistical genetic and clinical information for missense variants assayed in this study. Genomic coordinates correspond to the GRCh38 assembly. Predictions of clinical significance are taken from ClinVar (accessed on 6 August 2024). Allele frequencies and numbers of hemizygotes were obtained from gnomAD v4.1.0 (accessed on 6 August 2024) (1).

| **DNA Mutation** | **SNP ID** | **Protein Consequence** | **Clinical Significance** | **Allele Frequency (Hemizygotes)** |
| --- | --- | --- | --- | --- |
| g.23334970C > G |  | p.Pro32Arg | Likely pathogenic | 8.27 x 10^-7^ (0) |
| g.23335027G > A | -- | p.Ser51Asn | VUS | 9.11 x 10^-7^ (0) |
| g.23335092C > T | rs373105249 | p.Leu73Phe | Conflicting Classifications | 6.65 x 10^-5^ (21) |
| g.23335099C > A | -- | p.Pro75Gln | VUS | 0 (0) |
| g.23335099C > T | rs1198691680 | p.Pro75Leu | VUS | 8.94 x 10^-6^ (1) |
| g.23335180A > G | rs138264321 | p.Gln102Arg | VUS | 7.29 x 10^-6^ (4) |
| g.23379687G > A | rs747642714 | p.Val150Met | VUS | 7.44 x 10^-6^ (3) |
| g.23379781A > C | rs1175938564 | p.Lys181Thr | Likely Pathogenic | 9.11 x 10^-7^ (1) |
| g.23379822G > A | rs769407241 | p.Val195Ile | VUS | 5.65 x 10^-5^ (25) |
| g.23379877A > G | -- | p.Tyr213Cys | Conflicting Classifications | 0 (0) |
| g.23380146G > A | rs1060499615 | p.Gly303Arg | VUS | 0 (0) |
| g.23393099C > A | -- | p.Asp527Glu | VUS | 1.91 x 10^-5^ (11) |
| g.23393164T > G | -- | p.Phe549Cys | VUS | 0 (0) |
| g.23393322A > G | -- | p.Thr602Ala | VUS | 1.82 x 10^-6^ (0) |

**Table S3: *In silico* predictions for pathogenicty of PTCHD1 missense variants.**

In order to predict the consequences of point mutations on protein pathogenicity, ten separate computational algorithms were employed: PROVEAN (2), SIFT (3), PPH2 (4), Condel (5), CADD (6), REVEL (7), MetaLR (8), Mutation Assessor (9), InMeRF (10), and MPC (11). Missense variants were deemed to be likely pathogenic or benign based on algorithm-specific thresholds for pathogenicity. Red and green cells indicate that the missense mutation is predicted to be likely pathogenic or benign, respectively, according to the pathogenicity threshold for the given algorithm. For the PPH2 algorithm, yellow cells indicate possible predicted missense variant pathogenicity.

| **Missense Variant** | **PROVEAN** | **SIFT** | **PPH2** | **Condel** | **CADD** | **REVEL** | **MetaLR** | **Mutation Assessor** | **InMeRF** | **MPC** |
| --- | --- | --- | --- | --- | --- | --- | --- | --- | --- | --- |
| p.Ser51Asn | 0.03191 | 0.84 | 0 | 0.42475 | 21.2 | 0.25400 | 0.32900 | 0.08100 | 0.28100 | 0.50897 |
| p.Leu73Phe | 0.03041 | 0.13 | 0.127 | 0.54109 | 22.7 | 0.55500 | 0.57800 | 0.52800 | 0.56500 | 0.89012 |
| p.Pro75Gln | 0.90332 | 0.03 | 0.978 | 0.55643 | 25.9 | 0.84300 | 0.56000 | 0.56000 | 0.91700 | 0.88235 |
| p.Pro75Leu | 0.78046 | 0 | 1 | 0.55652 | 28.6 | 0.83700 | 0.33700 | 0.56000 | 0.56000 | 0.86978 |
| p.Gln102Arg | 0.17834 | 0.61 | 0 | 0.38084 | 18.6 | 0.26200 | 0.02100 | 0.00300 | 0.23100 | 0.56322 |
| p.Val150Met | 0.19933 | 0.04 | 0.236 | 0.53861 | 23.3 | 0.39500 | 0.69600 | 0.33800 | 0.59900 | 0.76398 |
| p.Lys181Thr | 0.14193 | 0.19 | 0.001 | 0.53828 | 21.9 | 0.28600 | 0.49000 | 0.40300 | 0.48200 | 0.61421 |
| p.Val195Ile | 0.20791 | 0.08 | 0.709 | 0.54120 | 21.8 | 0.62200 | 0.74600 | 0.37800 | 0.92200 | 0.63626 |
| p.Tyr213Cys | 0.96395 | 0 | 0.995 | 0.53725 | 26.5 | 0.91500 | 0.77800 | 0.30200 | 0.78800 | 0.89993 |
| p.Asp527Glu | 0.15578 | 0.15 | 0.752 | 0.53206 | 23.3 | 0.52000 | 0.63200 | 0.41100 | 0.85600 | 0.87029 |
| p.Phe549Cys | 0.94899 | 0 | 1 | 0.56124 | 28.0 | 0.91400 | 0.75500 | 0.56000 | 0.88200 | 0.92409 |
| p.Thr602Ala | 0.06026 | 1 | 0.996 | 0.48406 | 20.9 | 0.47800 | 0.35800 | 0.16000 | 0.86300 | 0.87650 |
